# Shear Relaxation Governs Dynamic Processes of Biomolecular Condensates

**DOI:** 10.1101/2021.04.17.440275

**Authors:** Archishman Ghosh, Divya Kota, Huan-Xiang Zhou

## Abstract

Phase-separated biomolecular condensates must respond agilely to biochemical and environmental cues in performing their wide-ranging cellular functions, but our understanding of condensate dynamics is lagging. Ample evidence now indicates biomolecular condensates as viscoelastic fluids, where shear stress relaxes at a finite rate, not instantaneously as in viscous liquids. Yet the fusion dynamics of condensate droplets has only been modeled based on viscous liquids, with fusion time given by the viscocapillary ratio (viscosity over interfacial tension). Here we used optically trapped polystyrene beads to measure the viscous and elastic moduli and the interfacial tensions of four types of droplets. Our results challenge the viscocapillary model, and reveal that the relaxation of shear stress governs fusion and other dynamic processes of condensates.

---

Phase-separated biomolecular condensates mediate cellular functions ranging from stress response to chromatin organization ^1–3^. Condensates, including membraneless organelles such as stress granules ^1, 2^, P granules ^4, 5^, and nucleoli ^6^, typically appear as micro-sized droplets and span material states from liquid-like to solid-like, or become solid-like over time (a process known as maturation or aging ^7–11^). In stress granules and many other cases, a dynamic, liquid state allows for rapid assembly, disassembly, or clearance in response to biochemical or environmental cues and for easy exchange of ligands or macromolecular components with the surrounding bulk phase ^1, 2, 4, 12–14^. Condensate solidification has been implicated in neurodegeneration and other pathologies ^1, 7, 9, 14, 15^. Thus an ATP-dependent process actively maintains the dynamics of nucleoli ^12^, and RNA and chaperones prevent condensates from solidification ^1, 16, 17^. In other cases, the extent of liquidity is tuned either temporally or spatially for appropriate condensate assembly or function ^5, 8, 18, 19^. Despite the crucial importance of condensate dynamics, its quantification and relation to molecular properties are still elusive.

Liquid droplets have a tendency to fuse and relax into a spherical shape, and the fusion speed has been used as an indicator of condensate dynamics ^4, 6, 7, 9, 12, 18, 20–27^. All fusion data have been analyzed by modeling condensates as purely viscous (i.e., Newtonian) liquids, where fusion, driven interfacial tension (capillarity; *γ*) but retarded by viscosity (*η*), occurs on the viscocapillary timescale *τ*_vc_ = *ηR*/*γ*, where *R* denotes droplet radius. This viscocapillary model based on Newtonian fluids has not been tested by measuring fusion speed, viscosity, and interfacial tension at the same time. Recent theoretical calculations have cast doubt on its validity if condensates are viscoelastic, i.e., partly liquid and partly solid ^28^. A crucial difference between viscous liquids and elastic solids lies in the shear relaxation modulus, *G*(*t*), which measures how shear stress relaxes upon the introduction of a unit-step strain at time *t* = 0 (see Supplementary Text). In many ways *G*(*t*) is similar to the memory kernel of a generalized Langevin particle. In viscous liquids, shear relaxation is instantaneous and *G*(*t*) is a delta function of time, *ηδ*(*t*). In elastic solids, shear stress never relaxes and hence *G*(*t*) is a constant. In viscoelastic fluids, shear relaxation occurs at a finite rate, as exemplified by the Maxwell fluid, with *G*(*t*) given by an exponentially decaying function of time (Supplementary Fig. 1). The Fourier transform of *G*(*t*) defines the complex shear modulus *G*\*(*ω*), whose real and imaginary parts are the elastic and viscous moduli, respectively. Here *ω* denotes the angular frequency. *G*\*(*ω*) data have been reported for some condensates, indicating that they are indeed viscoelastic on the ms to s timescales ^10, 29–32^. To assess the viscocapillary model and to uncover other determinants of the fusion speed, here we used optical tweezers (OT) to measure the complex shear moduli and the interfacial tensions of diverse condensates. The fusion speeds of these condensates were reported previously ^26^. The macromolecular components (Fig. 1A) include the single-domain protein lysozyme (L), pentameric constructs of SH3 domains (S) and SH3-targeting protein-rich motifs (P), and two polymers: polylysine (pK) and heparin (H). L, P, and pK carry significant net positive charges whereas S and H carry significant negative charges. Oppositely charged, binary mixtures comprising pK:H, p:H, S:P, and S:L form droplets, which fall under gravity, fuse, and settle on a coverslip with tallness sustained by interfacial tension (Fig. 1B).

**Fig. 1.**
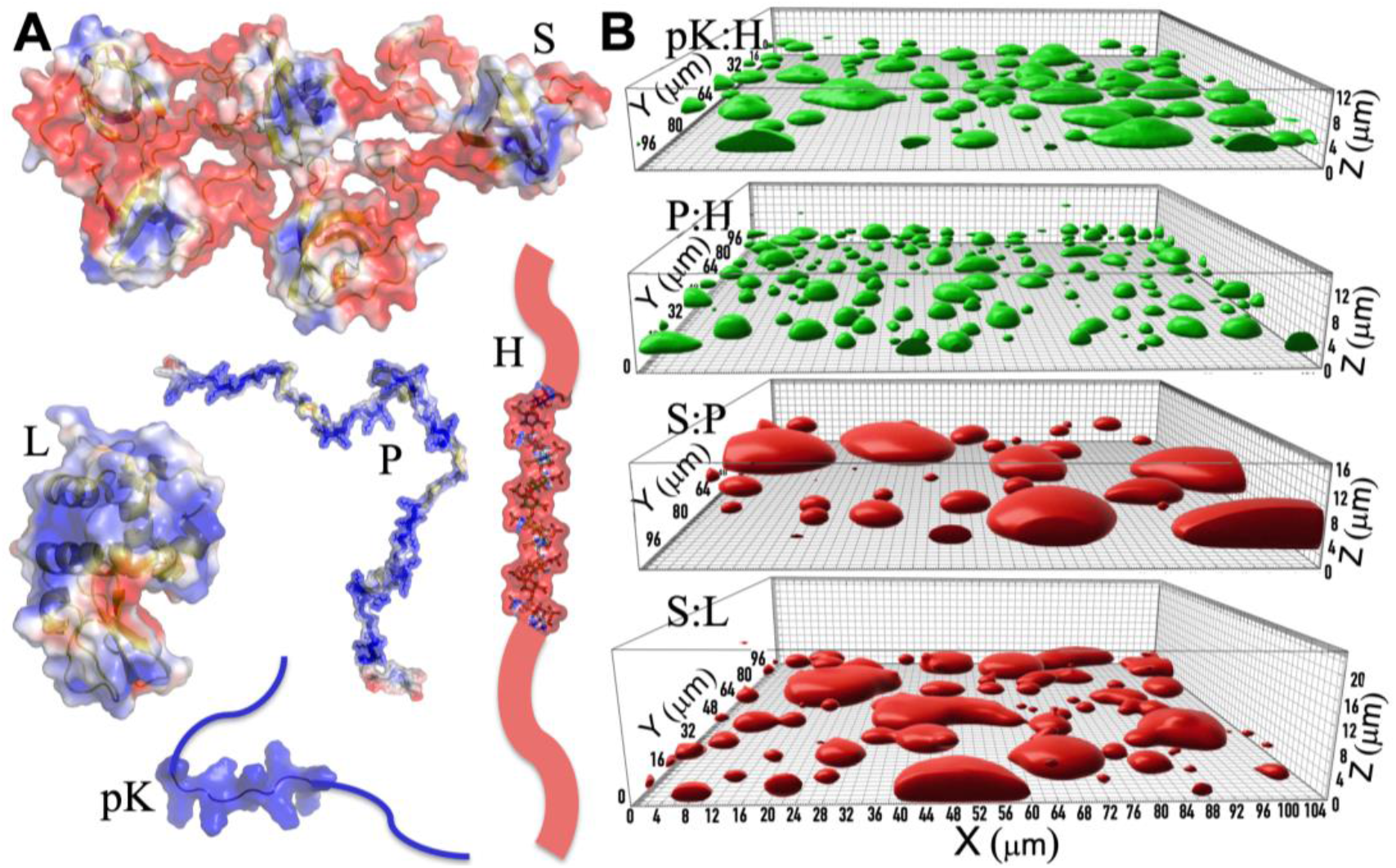
Macromolecular components and the resulting four types of droplets. (**A**) Molecular structures of pentameric constructs of SH3 domains (S) and proline-rich motifs (P), lysozyme (L), heparin (H), and polylysine (pK), rendered by electrostatic surfaces (blue and red: positive and negative electrostatic potentials, respectively). (**B**) pK:H, P:H, S:P, and S:L droplets settled on a coverslip, visualized by the fluorescence of either FITC-labeled H (green) or Alexa 594-labeled S (red) reproduced from ^26^. Growing tallness of droplets in the series gives crude indication of a modest increase in interfacial tension from pK:H to S:L.

## Results

We probed the viscoelasticity inside the settled droplets by oscillating an optically trapped polystyrene bead ^33, 34^ (Figs. 2A,B; see Supplementary Information for details and Movie S1 for illustration). The resulting shear relaxation moduli fit well to a combination of two Maxwell components (Fig. 3A-D),

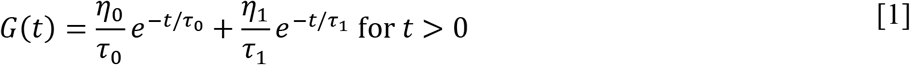

which is known as the Burgers model for linear viscoelasticity. *τ*_0_ and *τ*_1_ are relaxation times, and the corresponding amplitudes, *η*_0_ and *η*_1_, are viscosities. The resulting elastic and viscous moduli are

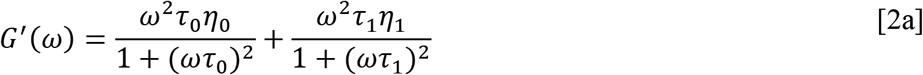

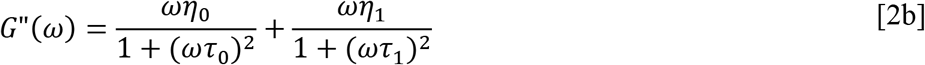

**Fig. 2.**
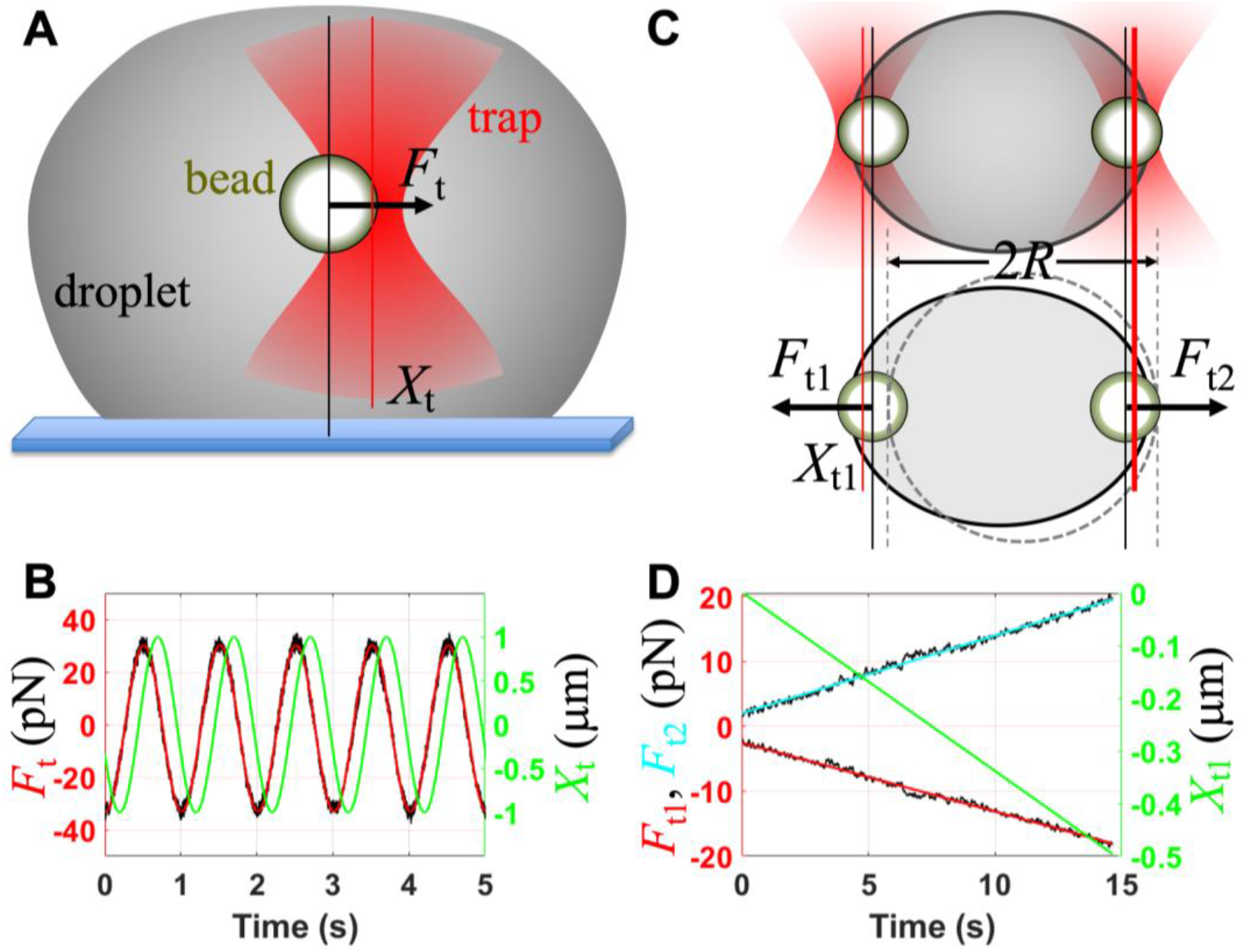
Design of experiments for measuring viscous and elastic moduli and interfacial tension. (**A**) Oscillatory micro-rheology inside a settled droplet. (**B**) Time traces of the trap position (*X*_t_; green) and trapping force (*F*_t_; black) on a bead inside a pK:H droplet. A fit of the force trace to a cosine function of time is shown in red. The oscillation frequency (*ω/2π*) was 1 Hz. (**C**) A droplet suspended by two trapped beads at the opposite poles (top: side view; bottom, top view). Trap 2 was fixed in place while trap 1 was moved toward the left. (**D**) Traces of the trap 1 position (*X*_t1_; green) and traps 1 and 2 forces (*F*_t1_ and *F*_t2_; black) measured on a pK:H droplet. The force traces were smoothed by moving average over a 64-ms window; linear fits are overlaid. Trap 1 was moved at a constant speed of approximately 0.05 μm/s.

**Fig. 3.**
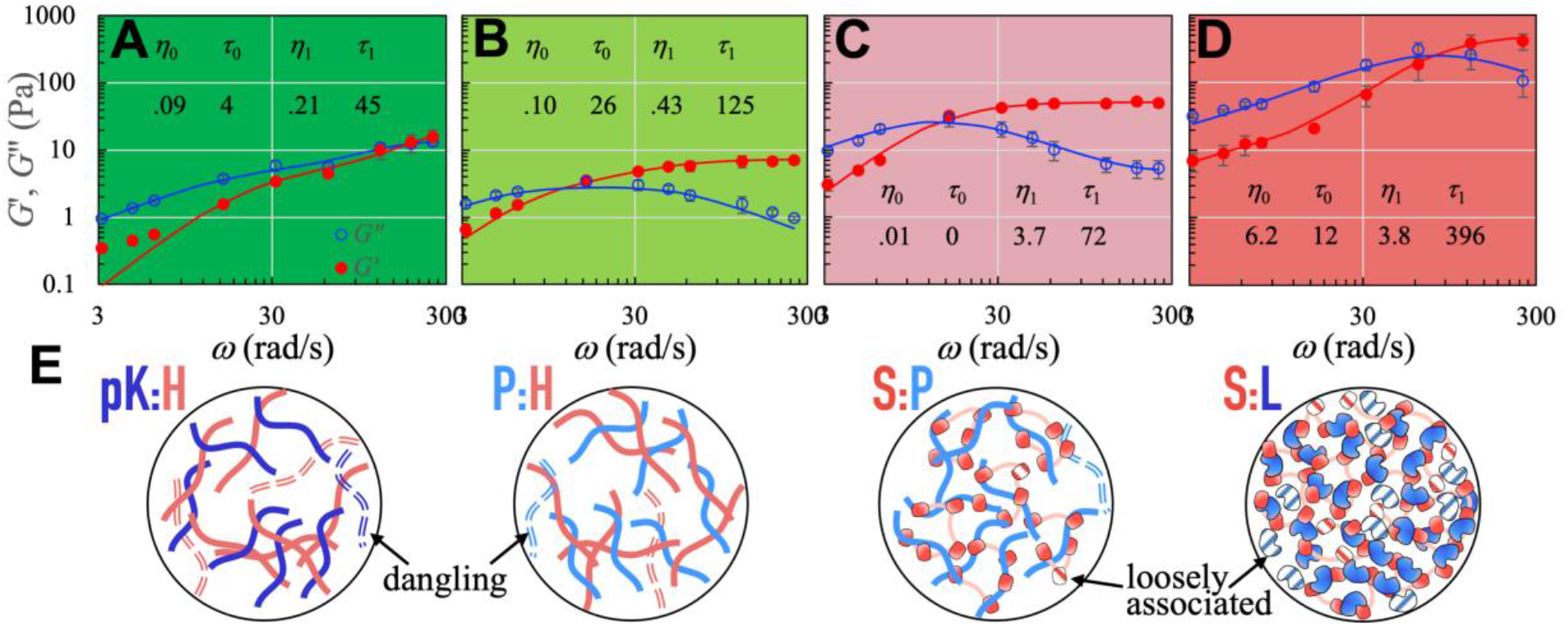
Complex shear moduli and macromolecular networks of four types of droplets. (**A-D**) Viscous (*G*′) and elastic (*G*″) moduli of pK:H, P:H, S:P, and S:L droplets. Red filled circles and blue open circles display *G*′ and *G*″, respectively; error bars display standard error of the mean among replicate measurements. Fits of the data to the Burgers model (Eqs [2a,b]) are shown as red and blue curves. The fitting parameters are listed in the figures. (**E**) Illustration of macromolecular networks inside droplets. Dangling polymer chains are shown as dash; loosely associated domains are shown as hatched fill.

According to transient network models for associative polymers, the shorter time constant, *τ*_0_, is dictated by the conformational dynamics of the macromolecular components, whereas the longer time constant, *τ*_1_, is dictated by the kinetics of macromolecular transient association and dissociation ^35, 36^. The amplitude ratio, *η*_0_/*η*_1_, is affected by the population ratio of dangling or loosely associated macromolecules to tightly associated ones (Fig. 3E).

In S:P droplets, *τ*_0_ is 0, which means that the first component of the shear relaxation modulus is actually Newtonian. In the other three types of condensates, *τ*_0_ is 5- to 35-fold shorter than the corresponding *τ*_1_. Most condensate dynamic processes of interest (e.g., droplet fusion) occur on timescales longer than *τ*_0_; then reconfiguration of macromolecular networks inside condensates, with a time constant *τ*_1_, is the main mode of shear relaxation. Although the inverse of the crossover angular frequency *ω*_x_ [defined by *G*′(*ω_x_*) = *G*″(*ω*_x_)] has commonly been identified as the network reconfiguration time (e.g., ref ^36^), *ω*_x_ for the Burgers model actually depends on *η*_0_/*η_1_* (Supplementary Fig. 2). 1/*ω*_x_ is close to *τ*_1_ only when *η*_0_/*η*_1_ is < 0.26. This condition is not satisfied in two (pK:H and S:L) of the four types of condensates studied, where 1/*ω*_x_ is closer to *τ*_0_ instead of *τ*_1_. S:L is the only case where *η*_0_/*η*_1_ is > 1, likely due to a high level of loosely associated L molecules (Fig. 3E). In all the four types of condensates, the network reconfiguration time *τ*_1_ is tens to hundreds ms.

In dynamic processes that are much slower than macromolecular network reconfiguration, shear relaxation is then fast and condensates behave entirely as Newtonian fluids, with viscosity *η* = *η*_0_ + *η*_1_. Fluorescence recovery after photobleaching (FRAP) is such an example, with time constants (*τ*_FR_) ranging from 2.1 ± 0.2 s to 105.1 ± 2.3 s for pK:H, P:H, S:P, and S:L condensates ^26^. Other methods (e.g., particle tracking) that work on similar or longer timescales would also only reveal Newtonian behaviors ^6, 20, 21, 27, 37–39^. In a Newtonian fluid, *τ*_FR_ is inversely proportional to the diffusion constant of the fluorescently labeled species and proportional to the fluid viscosity. These measured *τ*_FR_ values do correlate with the zero-shear rate viscosities (*η*) measured here by OT-based oscillatory micro-rheology, which are 0.30 ± 0.03, 0.53 ± 0.04, 3.75 ± 0.14, and 10.1 ± 1.1 Pa s for the four types of condensates. Indeed, the viscosities deduced from *τ*_FR_ (see Supplementary text) show quantitative agreement with the OT-measured values (Fig. 4A). The viscosity of S:L condensates is 33 times higher than that of pK:H condensates. In reference to the viscosity of water (*η*_w_; = 8.9 × 10^−4^ Pa s at 25 °C), the relative viscosities (*η*/*η*_w_) of the four types of condensates are 340, 600, 4200, and 11000.

**Fig. 4.**
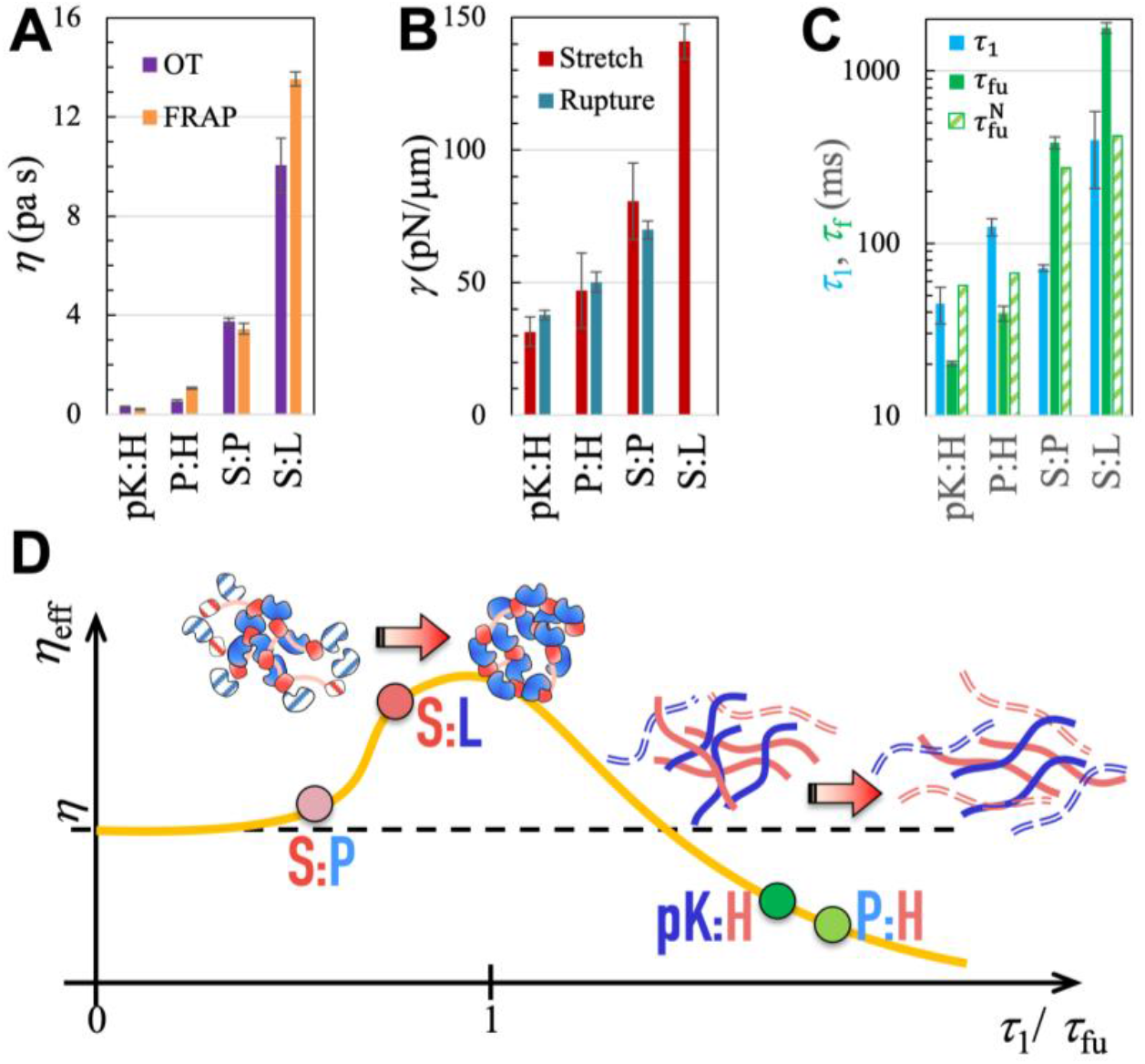
Deviations of condensates from Newtonian fluids. (**A**) Comparison of zero-shear viscosities from OT measurements and deduced from FRAP time constants. Error bars represent errors of parameters in fitting data to models. For S:L droplets, the L concentration was 300 μM in OT measurements but 2000 μM in FRAP; a somewhat higher *η* is thus expected in the FRAP experiment. (**B**) Interfacial tensions from OT measurements, by either stretching or rupturing droplets. For S:L droplets, the rupture force exceeded the trapping power of the instrument and hence only a lower bound of 100 pN/μm could be placed for *γ*. (**C**) Comparison of shear relaxation time (*τ*_1_), measured fusion (*τ*_fu_), and predicted fusion time 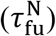 by the viscocapillary model. (**D**) Shear thickening at *τ*_1_/*τ*_fu_ close to 1 and shear thinning at longer *τ*_1_/*τ*_fu_. The solid curve displays characteristic behaviors of associative polymers. The illustration at top shows shear-induced strengthening of macromolecular networks; the illustration at right shows the opposite effect.

Two main determinants of viscosity are macromolecular interaction strength and degree of macromolecular structural compactness ^26^. Between pK:H and P:H, the slightly lower viscosity of the former can be attributed to a higher salt concentration (i.e., 1.0 M KCl, which enabled pK:H to form liquid droplets; at the 0.15 M KCl concentration in the other three types of condensates, pK:H only formed solid-like aggregates). KCl decreases viscosity by screening electrostatic attraction between the oppositely charged components. When the KCl concentration in P:H condensates is increased from 0.15 M to 0.3 and 0.4 M, the zero-shear viscosity decreases from 0.53 ± 0.04 Pa s to 0.25 ± 0.02 Pa s and 0.22 ± 0.01 Pa s, respectively. The components of pK:H and P:H condensates are polymers, which loosely pack but are connected to each other by numerous inter-chain bridges (Fig. 3E, left two panels). In contrast, the components of S:L condensates are a single folded domain and a pentamer of folded domains; inter-domain association results in close macromolecular packing (Fig. 3E, right most panel). The increase in viscosity going from P:H to S:P to S:L thus largely reflects the rise in the structural compactness of the components, leading to higher packing densities. The relative viscosity of a pure L solution at a high concentration of 480 g/L (or 34 mM) is only ~270 at 25 °C ^40^. The relative viscosity *η*_0_/*η*_w_ attributable to loosely associated L in S:L condensates is 7000. The latter much greater value means that even those loosely associated L molecules have significant interactions with surrounding macromolecular networks. In the pure L solution, a decrease in temperature to 5 °C effectively strengthened the attraction among L molecules and increased the relative viscosity to 2000 ^40^.

Let us now compare the network reconfiguration times (*τ*_1_) among the four types of condensates. pK:H condensates have the shortest *τ*_1_ at 45 ± 11 ms. One reason again is the high concentration of KCl, which weakens inter-chain electrostatic interactions and thereby inter-chains bridges more readily break. This conclusion is supported by the decrease in *τ*_1_ in P:H condensates, from 125 ± 14 ms to 56 ± 10 ms and 36 ± 3 ms, respectively, when the KCl concentration is deceased from 0.15 M to 0.3 and 0.4 M. S:L condensates have the longest *τ*_1_, at 396 ± 19 ms. As noted previously ^26^, breaking the extensive contacts between folded domains takes more energy and hence occurs more slowly than breaking bridges between polymer chains. S:P condensates have a *τ*_1_ (72 ± 3 ms) that is shorter than the counterparts in both P:H and S:L condensates. Of the two components of S:P condensates, one is a polymer and the other is a pentamer of folded domains. The resulting macromolecular networks are neither as entangled as in P:H condensates nor as densely packed as in S:L condensates, therefore explaining the shorter *τ*_1_. We note that in condensates formed by quaternary mixtures of S, P, H, and L, the components demix to generate P:H-rich and S:L-rich foci ^41^. This phenomenon is probably related to the rapid breakup of S:P networks as indicated by the short shear relaxation time *τ*_1_.

We measured the interfacial tensions (*γ*) of pK:H, P:H, S:P, and S:L droplets by stretching them using two trapped beads positioned at the opposite poles ^30^ (Figs. 2C,D; see Supplementary Information for details and Movie S2 for illustration). The values of *γ* are 31.5 ± 5.5, 47.0 ± 14.1, 80.7 ± 14.3, and 140.9 ± 6.6 pN/μm, respectively (displayed in Fig. 4B). They follow the same order as the zero-shear viscosities, and reflect successive strengthening of macromolecular networks. The ratio of interfacial tensions between the last member (S:L) and the first member (pK:H) of the series is 4.5. The droplet stretching method for interfacial tension was validated by measuring the force required to rupture the droplet surface when a trapped bead was pulled from inside (Fig. 4B; Movie S3 and Supplementary Fig. 3).

With both *η* and *γ* at hand, we can predict the fusion time of two equal-sized droplets (with radius *R*) if condensates inside are Newtonian fluids. This is given by ^26^

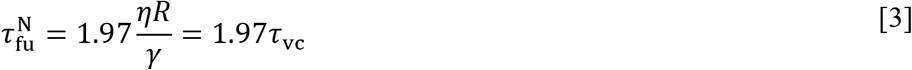

Because the zero-shear viscosities of the four types of droplets vary by 33-fold while the interfacial tensions vary by 4.5-fold, the viscocapillary model predicts an order of magnitude difference in fusion time between S:L and pK:H droplets. However, the measured fusion times (*τ*_fu_), reported previously ^26^, vary by close to two orders of magnitude among the four types of droplets. The measured and predicted fusion times are compared individually in Fig. 4C. The prediction is close for S:P, but is too high for pK:H and P:H whereas too low for S:L. The discrepancies of 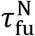 from *τ*_fu_ are in opposite directions for the first and last members of the series, which account for the much narrower range of the predicted fusion times.

## Discussion

It is thus clear that, whereas S:P condensates behave as Newtonian fluids during droplet fusion, the other three types of condensates do not. S:L condensates deviate from Newtonian fluids in one direction 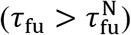 but pK:H and P:H condensates deviate from Newtonian fluids in the opposite direction 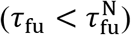. The opposite non-Newtonian behaviors indicate shear thickening (increase in viscosity) and shear thinning (decrease in viscosity), respectively. At increasing shear rate (i.e., rate of relative deformation; denoted by 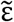), associative polymer solutions generally exhibit three regimes: Newtonian behavior at low 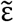; shear thickening at 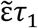 around 1; and shear thinning at high 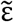 (e.g., ref ^36^). For droplet fusion, 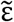 is measured by 1/*τ*_fu_ (relative deformation of order 1 during time *τ*_fu_). We thus expect the effective viscosity (*η*_eff_) to have a dependence on *τ*_1_/*τ*_fu_ that is characteristic of associative polymer solutions (orange curve in Fig. 4D). S:P condensates behave as Newtonian fluids because their *τ*_1_ is short relative to the fusion time. S:L condensates have a longer *τ*_1_ and thus exhibit shear thickening, which arises due to shear-induced conversion of loose association into close association (Fig. 4D, illustration at top). pK:H and P:H condensates have reconfiguration times longer than their respective fusion times, and thus exhibit shear thinning, due to shear-induced breakup of inter-chain bridges (Fig. 4D, illustration at right).

We have demonstrated that diverse condensates are viscoelastic on the ms to s timescales. During fusion, condensates can deviate from Newtonian fluids in opposite directions (i.e., shear thickening or thinning), depending on the relative rates of shear relaxation and condensate deformation. The main mode of shear relaxation inside condensates is reconfiguration of macromolecular networks. While determination of phase diagrams yields information on energetic properties of macromolecular components, measurement of shear relaxation moduli allows us to probe dynamic properties of these molecules. Similar to fusion, other condensate dynamic processes, including shape recovery after deformation, physical association of different condensates, multiphase organization, assembly and disassembly, and aging, all involve reconfiguration of macromolecular networks. Shear relaxation provides the governing measure of condensate dynamics.

## Methods

Materials, methods and data analysis are described in detail in the Supplementary Information.

## Author contributions

Conceptualization: AG, HXZ

Methodology: AG, DK

Investigation: AG, DK, HXZ

Funding acquisition: HXZ

Project administration: HXZ

Supervision: HXZ

Writing: HXZ

## Data and materials availability

All data and codes are available from the corresponding author

## Supplementary information

Materials and methods

Supplementary text

Figs. S1 to S3

Supplementary references (*42–47*)

