## Supporting Text and Figures for "Shear Relaxation Governs Dynamic Processes of Biomolecular Condensates"

**This PDF file includes:**

Materials and methods
Supplementary text
Figs. S1 to S3
Supplementary references (*42–47*)
Captions for Movies S1 to S3

**Other Supplementary Materials for this manuscript include the following:**

Movies S1 to S3

### Materials and Methods

#### *Sample preparation*

Preparation of biomolecular droplets containing polystyrene beads for optical trapping was modified from a procedure in a previous study on fusion dynamics (*26*). Pentameric constructs of SH3 domains (S) and proline-rich motifs (P) were expressed and purified as reported, as was the labeling of S to produce Alexa 594-S [42]. The sources for lysozyme (L), heparin (H), FITC-heparin, polylysine (pK), and Ficoll70 were the same as reported [41, 42]. Carboxylate-coated polystyrene beads (2 µm diameter) were procured from Polysciences Inc. (Catalog# 18327-10). The working buffer was 10 mM imidazole pH 7 with 0.01% (w/v) $NaN_3$ and KCl at 0.15 M unless otherwise indicated. Droplet samples containing beads were prepared by sequentially adding buffer, KCl if needed, pre-dissolved Ficoll70, pre-diluted bead solution, the negatively charged macromolecular component, and finally the positively charged component. The final macromolecular concentrations (in µM) were: pK:H = 50:50; P:H = 40:40; S:P = 40:40; and S:L = 20:300; the Ficoll70 concentration was 50 g/L. For pK:H droplets, the KCl concentration was 1.0 M; P:H droplets were prepared at 0.15 M, 0.3 M, and 0.4 M KCl. These samples matched in composition with those the study on fusion dynamics (*26*), except for the presence of beads, which were at a concentration of approximately 0.01% (w/v). All sample preparations and optical tweezer (OT) based experiments were done at room temperature.

#### *Calibration of optical trap stiffness*

The stiffnesses of the traps in a LUMICKS C-Trap™ dual-trap OT instrument were calibrated by fitting the power spectrum of a trapped bead's position to that of a Brownian harmonic oscillator. Bead samples for calibration were diluted to a concentration of ~0.01 % (w/v) in deionized water. Stiffness calibration was also carried out in glycerol solutions to validate that the stiffness obtained was independent of the medium surrounding the trapped bead.

#### *Oscillation of a trapped bead inside settled droplets*

Carboxylate-coated beads inside settled droplets were oscillated at amplitudes of 1.0 to 0.25 µm and frequencies ($\omega/2\pi$) of 0.5 to 40 Hz by the LUMICKS C-Trap™ instrument (Fig. 2A). Following the preparation of bead-containing droplets, a 7-8 µL aliquot was loaded onto a custom-made sample chamber (*26*), and droplets were allowed (for approximately 10 min) to settle on the coverslip. The power of the trapping laser (100% given to trap 1) was then turned up sufficiently to trap a bead inside a settled droplet and oscillate the bead at a chosen amplitude and frequency.

Only sufficiently large droplets were selected so that a centrally trapped bead would be at least several bead diameters away from any boundary. A brightfield image guided the positioning of the trapped bead at a transversely central location inside the droplet; further positioning at the droplet center along the longitudinal (i.e., $z$) direction was achieved when the trapped bead and the droplet equator were both in sharp focus. $z$ positions that were either too high or two low (such that the bead got close to the top or bottom boundary) resulted in in-phase trap position and force traces at the lowest frequency (i.e., 0.5 Hz). In contrast, a $z$ position away from droplet boundaries resulted in a phase difference of $> \pi/4$ at 0.5 Hz.

A complete set of data was collected on each trapped bead and exported as hdf files containing the trap 1 $x$ position traces in volts and the trap 1 force $x$ component traces in pN over a few s, at frequencies from 0.5 to 40 Hz. Replicates ($N$ = 3 or 4) were performed on different droplets from the same sample or from different samples.

#### ***Data analysis for bead oscillation***

An optically trapped bead inside a viscoelastic material experiences three forces (*34*, *35*): a viscous force proportional to the velocity $U$ of the bead, an elastic force proportional to the displacement $X$ of the bead, and the trapping force $F_\mathrm{t}$ proportional to the displacement, $X - X_\mathrm{t}$, between the bead and trap. Neglecting inertial effects, we have the following force-balance equation:

$$-\xi_\mathrm{m} U - \kappa_\mathrm{m} X - \kappa_\mathrm{t}(X - X_\mathrm{t}) = 0 \qquad \text{[M1]}$$

where $\xi_\mathrm{m}$ is the friction coefficient proportional to the viscous modulus, $\kappa_\mathrm{m}$ is the spring constant proportional to the elastic modulus, and $\kappa_\mathrm{t}$ is the stiffness of the optical trap. The trap is driven in a sinusoidal motion, and its position can be expressed as

$$X_\mathrm{t} = X_{\mathrm{t}0} e^{i(\omega t + \phi)} \qquad \text{[M2]}$$

where $\omega$ is the angular frequency; its division by $2\pi$ gives the frequency in Hz. The expression in complex variables is for convenience; its real part is the quantity of interest. The trapping force can likewise be expressed as

$$F_\mathrm{t} = -\kappa_\mathrm{t}(X - X_\mathrm{t}) = F_{\mathrm{t}0} e^{i(\omega t + \phi + \Delta)} \qquad \text{[M3]}$$

From the last two equations, we can find the bead position as

$$X = X_0 e^{i(\omega t + \phi - \delta)} \qquad \text{[M4]}$$

where

$$X_0 e^{-i\delta} = X_{\mathrm{t}0} - \frac{F_{\mathrm{t}0}}{\kappa_\mathrm{t}} e^{i\Delta} \qquad \text{[M5]}$$

Using Eqs [M3] and [M4] in Eq [M1], we find

$$(\kappa_\mathrm{m} + i\omega\xi_\mathrm{m}) X_0 e^{-i\delta} = F_{\mathrm{t}0} e^{i\Delta} \qquad \text{[M6]}$$

The quantity, $\kappa_\mathrm{m} + i\omega\xi_\mathrm{m}$, in the parentheses defines the complex shear modulus $G^*(\omega)$,

$$6\pi a G^*(\omega) = \kappa_\mathrm{m} + i\omega\xi_\mathrm{m} \qquad \text{[M7]}$$

where $a$ is the radius of the trapped bead (1 μm in our case). Rearranging Eq [M6], we find

$$G^*(\omega) \equiv G'(\omega) + iG''(\omega) = \frac{F_{\mathrm{t}0} e^{i\Delta}}{6\pi a X_0 e^{-i\delta}} = \frac{F_{\mathrm{t}0}}{6\pi a X_{\mathrm{t}0}} \frac{e^{i\Delta}}{1 - \Upsilon e^{i\Delta}} \qquad \text{[M8]}$$

where in the last step we have used Eq [M5], and $\Upsilon = F_{\mathrm{t}0}/\kappa_\mathrm{t} X_{\mathrm{t}0}$. The real and imaginary parts, $G'(\omega)$ and $G''(\omega)$, are the elastic and viscous moduli, respectively. The final expressions for them are

$$G'(\omega) = \frac{F_{\mathrm{t}0}}{6\pi a X_{\mathrm{t}0}} \frac{\cos\Delta - \Upsilon}{(\cos\Delta - \Upsilon)^2 + \sin^2\Delta} \qquad \text{[M9a]}$$

$$G''(\omega) = \frac{F_{\mathrm{t}0}}{6\pi a X_{\mathrm{t}0}} \frac{\sin\Delta}{(\cos\Delta - \Upsilon)^2 + \sin^2\Delta} \qquad \text{[M9b]}$$

The trap position and force profiles from the bead oscillation experiments were analyzed using MATLAB scripts. Both signals were fit to a cosine wave function (Fig. 2B):

$$X_\mathrm{t} = X_{\mathrm{t}0} \cos(\omega t + \phi) + B_1 \qquad \text{[M10]}$$

$$F_{\mathrm{t}} = F_{\mathrm{t0}} \cos(\omega t + \phi + \Delta) + B_2 \quad \text{[M11]}$$

The resulting amplitudes $X_{\mathrm{t0}}$ and $F_{\mathrm{t0}}$ and the phase difference $\Delta$, along with the trap stiffness $\kappa_{\mathrm{t}}$, were used in calculate the elastic and viscous moduli according to Eqs [M9a,b]. $X_{\mathrm{t0}}$ was converted to μm using a conversion factor from volt to μm, obtained by assuming that the trap position amplitude at 0.5 Hz was the preset value (e.g., 1.0 μm). The resulting $X_{\mathrm{t0}}$ values at the highest frequencies (close to and at 40 Hz) could be somewhat less than the preset values.

The foregoing experimental and data analysis protocols were validated by oscillating trapped beads (at frequencies between 0.5 and 10 Hz) in deionized water and glycerol solutions. This control experiment yielded the viscosity of the surrounding medium as $G''(\omega)/\omega$. The results were consistent with literature values.

***Stretching of droplets suspended by two trapped beads at the opposite poles***

Droplets were suspended by two trapped beads at the opposite poles, in a configuration introduced by Jawerth et al. (*30*) (Fig. 2C, top panel). As first proposed in (*31*), the droplet was stretched to obtain its static spring constant $\chi_0$, which is proportional to the interfacial tension. Stretching is a more convenient protocol than the oscillatory one used by Jawerth et al., because it specifically and solely probes the interfacial tension.

To prepare the suspended configuration, all droplets and beads were first allowed to settle on the coverslip. Two beads were then trapped and lifted from the bottom. Around each lifted bead, a droplet grew. By bringing the two beads toward each, the droplets fused spontaneously to generate the suspended configuration, where the trapped beads are located at the opposite poles of the fused droplet. In cases where settled beads were difficult to lift due to high viscosity or interfacial tension (as found in S:L samples), beads were trapped before settling on the coverslip in order to generate the suspended configuration; the suspended droplet was then held until all neighboring beads and droplets had settled. From this configuration, trap 2 was fixed while trap 1 was pulled away at a very low speed (0.05 μm/s), for a total of 0.5 μm. Pulling speeds of 0.01 μm/s and 0.1 μm/s were also tested to verify that these pulling speeds had no effect on the forces measured. Moreover, stretching and subsequent retracting yielded the same $\chi_0$, showing that the droplets indeed behaved like a spring. The data for trap 1 position and traps 1 and 2 force *x* components over the entire pulling period were exported as hdf files. The brightfield images of the entire process were also recorded in video. Replicates (*N* = 4 to 6) were performed on different suspended droplets.

***Data analysis for droplet stretching***

The static spring constant of a droplet is

$$\chi_0 = \frac{(F_{\mathrm{t2}} - F_{\mathrm{t1}})/2}{\Delta X} \quad \text{[M12]}$$

where $F_{\mathrm{t1}}$ and $F_{\mathrm{t2}}$ are the forces exerted by trap 1 and trap 2, respectively, at the opposite poles, and $\Delta X$ is the resulting stretch of the droplet. $\Delta X$ is measured by the change in the inter-bead distance. Instead of $\Delta X$, we directly tracked, $\Delta X_{\mathrm{t1}}$, the movement of trap 1 relative to trap 2, which was kept fixed (Fig. 2C, bottom panel). The quantity analogous to $\chi_0$ is

$$\chi_{\mathrm{sys0}} = \frac{(F_{\mathrm{t2}} - F_{\mathrm{t1}})/2}{-\Delta X_{\mathrm{t1}}} \quad \text{[M13]}$$

The trapping forces are

$$F_{tj} = -\kappa_{tj}(X_j - X_{tj}), \qquad j = 1 \text{ and } 2 \qquad \text{[M14]}$$

Force balance on the system in suspension yields

$$F_{t1} + F_{t2} = 0 \qquad \text{[M15]}$$

We can identify $\Delta X$ as $X_2 - X_1 - d_0$, where $d_0$ is the initial value of $X_2 - X_1$. Note $d_0$ is also the initial value of $X_{t2} - X_{t1}$ since initially $X_j = X_{tj}$. Moreover, $X_{t2}$ is held constant. Therefore $-\Delta X_{t1} = X_{t2} - X_{t1} - d_0$. Using these relations, we derive

$$\frac{1}{\chi_0} = \frac{1}{\chi_{sys0}} - \frac{1}{\kappa_{t1}} - \frac{1}{\kappa_{t2}} \qquad \text{[M16]}$$

This result reflects the fact that the system containing the droplet and two optical traps is equivalent to three springs connected in series.

We verified that, at the end of each stretching experiment, trap 1 indeed moved the preset amount (0.5 μm; converted from trap 1 position in volts using the aforementioned conversion factor). The trap 1 position changed as a linear function of time, though the negative of the slope, $v$, corresponding to the pulling speed, was somewhat less than the preset value (0.05 μm/s) as the pulling period was longer than the expected time of 10 s (see Fig. Fig. 2D). The trap 1 force *x* component and trap 2 force *x* component, after smoothing with moving average in a 64-ms window, were each fit to a linear function of time, with slopes $f_1$ and $f_2$. The static spring constant of the system comprising the droplet and two trapped beads was found as

$$\chi_{sys0} = \frac{(f_2 - f_1)/2}{v} \qquad \text{[M17]}$$

This result along with the stiffnesses of the traps were then used to obtain the static spring constant $\chi_0$ of the droplet itself.

As shown previously (*31*), $\chi_0$ is proportional to the interfacial tension $\gamma$ of the droplet:

$$\chi_0 = \frac{\pi\alpha(\theta_0)}{2}\gamma \qquad \text{[M18]}$$

where $\theta_0$ is the polar angle spanned by the trapped bead at each pole and is approximately $a/(R-a)$, where $R$ and $a$ are the radii of the unstretched droplet and the bead, respectively. For $\theta_0$ up to 0.5, $\alpha(\theta_0)$ is accurately given by

$$\frac{1}{\alpha(\theta_0)} = -0.5\ln\theta_0 + 0.34 \qquad \text{[M19]}$$

The interfacial tension was thus found as

$$\gamma = \frac{1}{\pi}[\ln(R/a - 1) + 0.68]\chi_0 \qquad \text{[M20]}$$

We determined the diameter ($2R$) of the suspended droplet by measuring the edge-to-edge distance of its image at the start of the recorded video using imageJ. The measurement was in pixels, which was converted to μm using the conversion factor of 0.0866 μm per pixel. Droplet diameters ranged from 7 to 18 μm.

***Surface rupture of settled droplets by a trapped bead***

To validate the interfacial tensions measured by stretching droplets, we pulled a trapped bead inside as settled droplet until the bead ruptured the droplet surface. Similar to a Wilhelmy plate [43] or a du Noüy ring [44], the trapped bead acted as a micro-tensiometer.

The initial setup was similar to what is described under "***Oscillation of a trapped bead inside settled droplets***", except this time the trapped bead was lifted to near the top of the droplet, to ensure that the resulting surface protrusion did not adhere to the coverslip. The bead was pulled at a low constant speed (0.1 μm/s) until it ruptured the droplet surface. The trace of the trapping force was exported.

Similar to the situation with a Wilhelmy plate or a du Noüy ring, the maximum trapping force $F_{\mathrm{rup}}$, corresponding to the moment where the droplet surface is ruptured by the bead, is related to the force due to the interfacial tension. This tension force can be estimated as the product of $\gamma$ and $2\pi a$, the circumference of the bead's equator. Additionally, the bead is pulled back by intermolecular molecules between the coating carboxylates and the droplet macromolecular components. A crude assumption is that the latter force is proportional to the tension force. Force balance thus yields

$$F_{\mathrm{rup}} = 2\pi a f \gamma \qquad \text{[M21]}$$

where $f$ is a numerical constant. The rupture forces for the four types of droplets studied here ranked in the same order as the interfacial tensions measured by the droplet stretching method. By choosing $f$ = 1.6, the interfacial tensions converted from rupture forces according to Eq [M21] are in quantitative agreement with the values from droplet stretching.

### Supplementary Text

#### Viscous liquids vs. elastic solids: distinction in shear relaxation

In a viscous liquid, a bead experiences a frictional force that is proportional to its velocity. In contrast, in an elastic solid, the bead experience a resistance that is proportional to its displacement. A corresponding contrast between the viscous liquid and elastic solid exists when these materials are deformed by shearing. The resulting stress, $\tilde{\tau}$, is proportional to the shear rate $\tilde{\varepsilon} = \frac{\partial}{\partial t}\tilde{\sigma}$ in the liquid but is proportional to the shear strain $\tilde{\sigma}$ itself in the solid.

Viscous liquids and elastic solids are opposite extremes of viscoelastic fluids. The latter materials, including biomolecular condensates, generally behave as partly liquid and partly solid. There the stress is determined by the entire history of the shear rate:

$$\tilde{\tau}(t) = \int_{-\infty}^{t} dt' G(t-t')\,\tilde{\varepsilon}(t') \qquad \text{[S1]}$$

The function $G(t)$ is called the shear relaxation modulus. This expression for the shear stress is similar in form and substance to the frictional force on a generalized Langevin particle; the counterpart of $G(t-t')$ is the memory kernel. Note that $G(t-t')$ must be 0 when $t' > t$ (such that future shear rate does not affect present stress); hence the upper limit of the integral can extend to $+\infty$. In a purely viscous liquid (also known as a Newtonian fluid), the stress is affected by the shear rate only at the present time, not any earlier time. That is,

$$\tilde{\tau}(t) = \eta\tilde{\varepsilon}(t) \qquad \text{[S2a]}$$

where $\eta$ is the viscosity. Here the shear relaxation modulus has no memory, as represented by a delta function

$$G(t) = \eta\delta(t) \qquad \text{[S2b]}$$

and hence shear relaxation is instantaneous. In contrast, the shear relaxation modulus of an elastic solid is a constant (denoted as $G_0$), meaning that the stress never relaxes. The result is the expected strain-stress relation

$$\tilde{\tau}(t) = G_0 \int_{-\infty}^{t} dt' \frac{\partial}{\partial t'} \tilde{\sigma}(t')$$

$$= G_0 \tilde{\sigma}(t) \qquad \text{[S3]}$$

Let us further illustrate with a unit-step strain introduced at time $t$ = 0 (Fig. S1A):

$$\tilde{\sigma}(t) = \Theta(t) \qquad \text{[S4]}$$

Noting the derivative of the Heaviside step function $\Theta(t)$ is a delta function, we find

$$\tilde{\varepsilon}(t) = \delta(t) \qquad \text{[S5]}$$

Substituting into Eq [S1], we have

$$\tilde{\tau}(t) = G(t) \qquad \text{[S6]}$$

The shear relaxation modulus thus represents the stress in response to a unit step strain introduced at time $t$ = 0. In the viscous liquid, the stress disappears after $t$ = 0 (Fig. S1B, top); i.e., shear relaxation is instantaneously as stated already. In the elastic solid, the stress, once generated at $t$ = 0, stays forever (Fig. S1C, top). Viscoelastic fluids fall in between, with shear relaxation occurring over a time period that is between 0 and infinity. An example is a Maxwell fluid, with $G(t)$ given by an exponential function of time (Fig. S1D, top). When a Newtonian component and a Maxwell component are added, one arrives at the Jeffreys model of linear viscoelasticity. The combination of two Maxwell components (with different time constants; see Eq. [1] in the main text) makes up the Burgers model.

Another type of shear strain of common interest is a sinusoidal function of time,

$$\tilde{\sigma}(t) = \tilde{\sigma}_0 e^{i\omega t} \qquad \text{[S7a]}$$

The corresponding shear rate is

$$\tilde{\varepsilon}(t) = i\omega \tilde{\sigma}_0 e^{i\omega t} \qquad \text{[S7b]}$$

and the stress is

$$\tilde{\tau}(t) = i\omega \tilde{\sigma}_0 e^{i\omega t} \int_{-\infty}^{+\infty} dt' G(t') e^{-i\omega t'} \equiv G^*(\omega) \tilde{\sigma}(t) \qquad \text{[S8]}$$

The last identity generalizes Eq [S3] and formally defines the complex shear modulus, $G^*(\omega)$. The latter is essentially the Fourier transform of the shear relaxation modulus,

$$G^*(\omega) = i\omega \int_{-\infty}^{+\infty} dt' G(t') e^{-i\omega t'} \qquad \text{[S9]}$$

For a viscous liquid (Eq [S2b]), we have

$$G^*(\omega) = i\omega\eta \qquad \text{[S10]}$$

which has only an imaginary part (Fig. S1B, bottom). On the other hand, for an elastic solid, by comparing Eqs [S3] and [S8], we find

$$G^*(\omega) = G_0 \quad [S11]$$

which has only a real part (Fig. S1C, bottom). In general, viscoelastic fluids have both real and imaginary parts,

$$G^*(\omega) = G'(\omega) + iG''(\omega) \quad [S12]$$

The real part is called the elastic (or storage) modulus, whereas the imaginary part is called the viscous (or loss) modulus. The elastic and viscous moduli of a Maxwell fluid are shown in (Fig. S1D, bottom.

**Comparison of condensate viscosities by OT and by FRAP**

In a previous study (*26*), we fit fluorescence recovery after photobleaching (FRAP) data to an exponential function

$$F(t) = F(\infty)[1 - e^{-t/\tau_{\mathrm{FR}}}] \quad [S13]$$

Here we use the resulting time constant ($\tau_{\mathrm{FR}}$) to deduce the viscosity inside condensates. According to Soumpasis [45], the half-time, $\tau_{1/2} = (\ln 2)\tau_{\mathrm{FR}}$, and the radius, $r_{\mathrm{B}}$, of the bleached region, can be used to find the diffusion constant of the fluorescently labeled species as

$$D = \frac{0.224 r_{\mathrm{B}}^2}{\tau_{1/2}} = \frac{0.224 r_{\mathrm{B}}^2}{(\ln 2)\tau_{\mathrm{FR}}} \quad [S14]$$

In the FRAP experiments, $r_{\mathrm{B}}$ was kept at 1.39 μm; $\tau_{\mathrm{FR}}$ was found to be 2.1 ± 0.2, 10.6 ± 0.4, 26.8 ± 1.6, and 105.1 ± 2.3 s, respectively for pK:H, P:H, S:P, and S:L condensates. The fluorescently labeled species was H in the first two cases and S in the last cases. The samples were otherwise the same as in the present study, except the L concentration in preparing the S:L samples were 2000 μM instead of 300 μM in the present work. The higher L concentration makes the condensates denser and hence more viscous. Thus the FRAP-derived viscosity for the S:L condensates should be somewhat higher than the corresponding OT-derived value.

To find the viscosity in condensates, we compare $D$ calculated from $\tau_{\mathrm{FR}}$ using Eq [S14] to the diffusion constant, $D_0$, obtained for H or S determined in water. The viscosity of the condensates is

$$\eta = \frac{D_0 \eta_{\mathrm{w}}}{D} = \frac{(\ln 2) D_0 \eta_{\mathrm{w}} \tau_{\mathrm{FR}}}{0.224 r_{\mathrm{B}}^2} \quad [S15]$$

where $\eta_{\mathrm{w}} = 8.9 \times 10^{-4}$ Pa s is the viscosity of water at 25 °C. For H, $D_0$ for a polymer fraction with molecular weight around 18 kD (as in our samples) was approximately 60 μm$^2$/s at 20 °C [46]. Using the Stokes-Einstein relation and the viscosities of water, $D_0$ for H at 25 °C can be found to be 69 μm$^2$/s. For S in water, we use a scaling relation between $D_0$ (in μm$^2$/s at 20 °C) and molecular weight ($M$, in Dalton), $10^4/D_0 = 3.48 M^{1/3} + 5.82$, derived for globular proteins [47], along with $M$ = 42849 Dalton to find $D_0$ = 78 μm$^2$/s. The latter translates into $D_0$ = 90 μm$^2$/s at 25 °C.

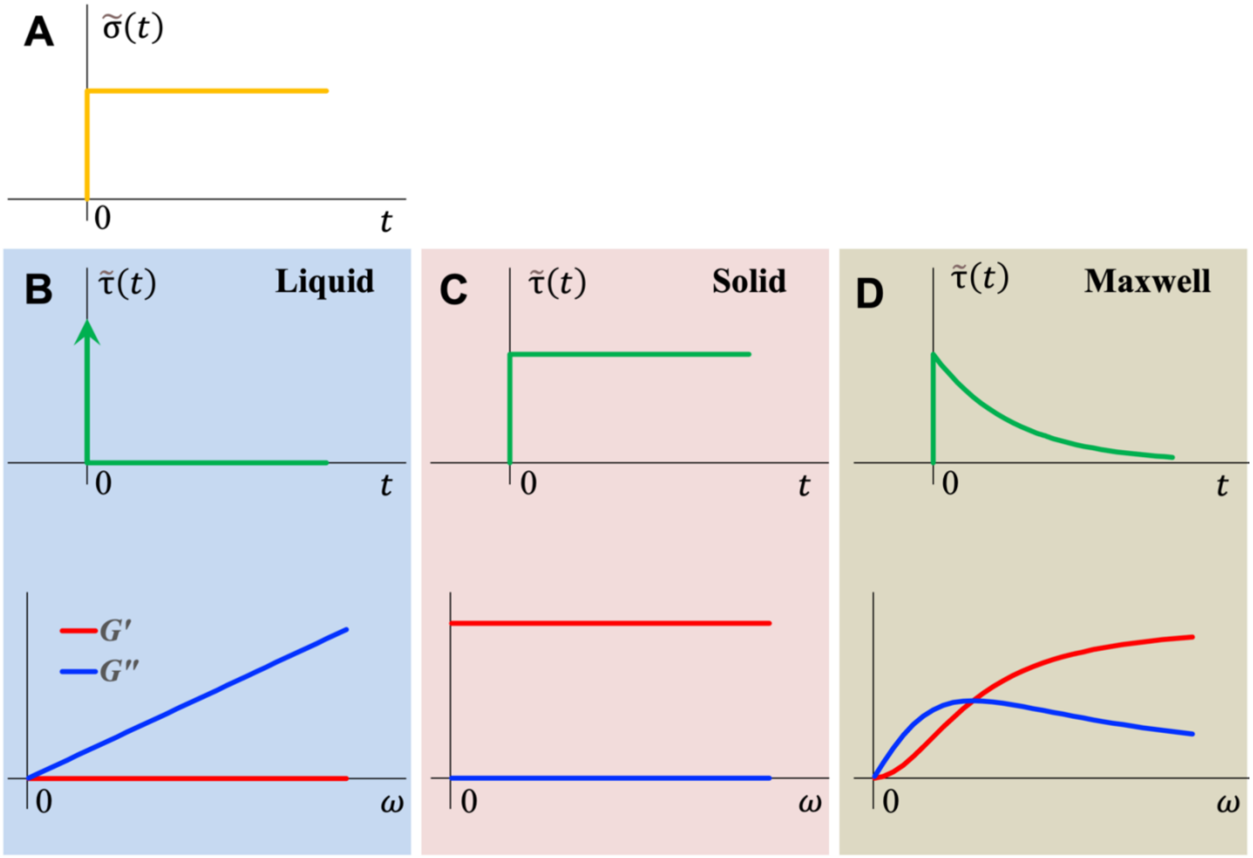


**Supplementary Fig. 1. Shear relaxation of liquids, solids, and viscoelastic fluids.** (**A**) A unit-step shear strain introduced at time $t = 0$. (**B-D**) Top: the resulting stress inside a viscous liquid, an elastic solid, or a Maxwell fluid. Bottom: the corresponding elastic and viscous moduli.

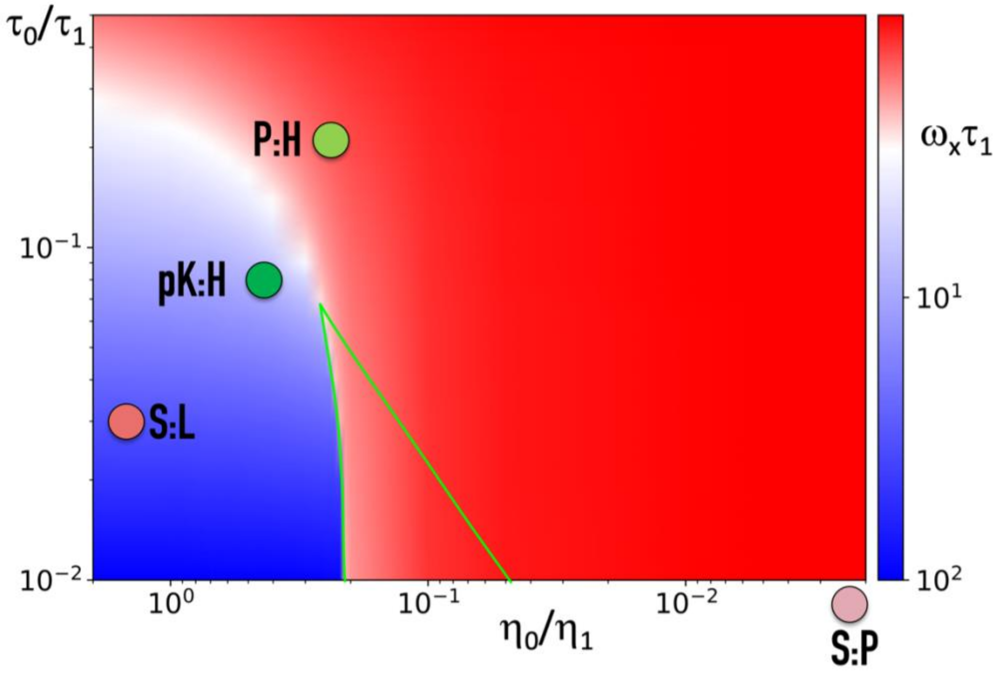


**Supplementary Fig. 2. Dependence of the crossover frequency $\omega_x$ on $\eta_0/\eta_1$ and $\tau_0/\tau_1$.** The value of $\omega_x\tau_1$ is displayed by color, according to the scale on the right. The pK:H, P:H, S:P, and S:L condensates are located on the map according to the measured $\eta_0/\eta_1$ and $\tau_0/\tau_1$; S:P has $\tau_0/\tau_1$ = 0 and is placed along the abscissa. Red regions have $\omega_x$ close to $1/\tau_1$ whereas blue regions have $\omega_x$ close to $1/\tau_0$. Inside the triangular region bordered by green curves, $G'(\omega)$ and $G''(\omega)$ cross each other three times; only the smallest crossover frequency is displayed.

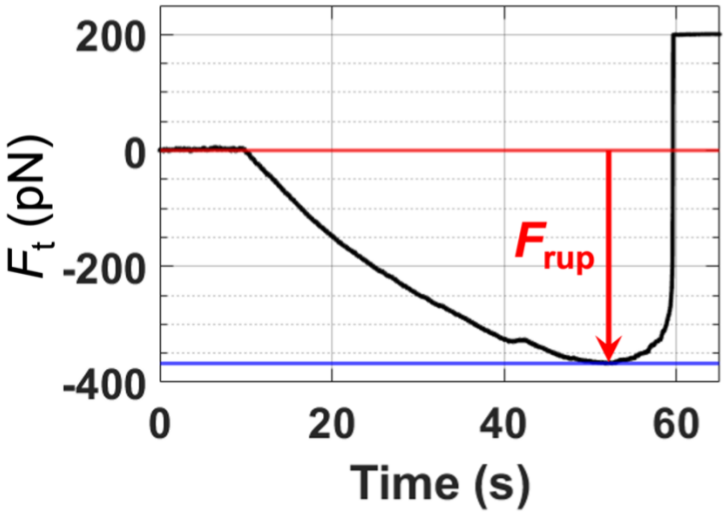


**Supplementary Fig. 3. Rupture force of a pK:H droplet.** The trap 1 force was smoothed by moving average over a 64-ms window. For this particular droplet, the rupture force was 367.9 pN, corresponding to an interfacial tension of 36.7 pN/μm according to Eq [M21] in Materials and Methods.

**Movie S1. Oscillating a trapped bead inside a settled pK:H droplet.** The trapped bead is seen as a white spot inside the droplet, oscillating at a frequency of 1 Hz.

**Movie S2. Stretching of a pK:H droplet suspended by two trapped beads at the opposite poles.** The trapped bead on the right is fixed in place while the trapped bead on the left is pulled toward the left, with a total displacement of 0.5 μm. The pulling speed is slightly less than 0.05 μm/s. The movie is played at four times the recording speed.

**Movie S3. Surface rupture of a pK:H droplet by pulling a trapped bead from inside.** The movie is played at eight times the recording speed.
